# Investigating and Assessing the Dermoepidermal Junction with Multiphoton Microscopy and Deep Learning

**DOI:** 10.1101/743054

**Authors:** Mikko J. Huttunen, Radu Hristu, Adrian Dumitru, Mariana Costache, Stefan G. Stanciu

## Abstract

Histopathological image analysis performed by a trained expert is currently regarded as the gold-standard in the case of many pathologies, including cancers. However, such approaches are laborious, time consuming and contain a risk for bias or human error. There is thus a clear need for faster, less intrusive and more accurate diagnostic solutions, requiring also minimal human intervention. Multiphoton Microscopy (MPM) can alleviate some of the drawbacks specific to traditional histopathology by exploiting various endogenous optical signals to provide virtual biopsies that reflect the architecture and composition of tissues, both in-vivo or ex-vivo. Here we show that MPM imaging of the dermoepidermal junction (DEJ) in unstained tissues provides useful cues for a histopathologist to identify the onset of non-melanoma skin cancers. Furthermore, we show that MPM images collected on the DEJ, besides being easy to interpret by a trained specialist, can be automatically classified into healthy and dysplastic classes with high precision using a Deep Learning method and existing pre-trained Convolutional Neural Networks. Our results suggest that Deep Learning enhanced MPM for in-vivo skin cancer screening could facilitate timely diagnosis and intervention, enabling thus more optimal therapeutic approaches.

## INTRODUCTION

As a result of their inherent optical sectioning capabilities and intrinsic contrast mechanisms, Multiphoton Microscopies (MPM) have emerged over the past three decades as very powerful tools for the label-free characterization of tissue morphology, functionality and biochemical composition (Hoover and Squier, 2013, König, 2018, Williams et al., 2001), in-vivo and ex-vivo. Among these, Two-Photon Excitation Fluorescence (TPEF) microscopy(So et al., 2000) and Second-harmonic Generation (SHG) microscopy(Campagnola and Loew, 2003), have demonstrated their usefulness for exploring important properties of tissues, which allow establishing their anatomical and functional states, and extracting valuable pathological cues(Balu et al., 2015, Muensterer et al., 2017, Zipfel et al., 2003a, Zipfel et al., 2003b)

TPEF involves the simultaneous absorption of two photons with combined energy sufficient to induce an electronic transition to an excited electronic state(So et al., 2000). Interestingly, TPEF allows to image in-vivo, ex-vivo or in-vitro the emission of various endogenous fluorophores such as NADH, FAD, melanin and others. Subsequently, TPEF microscopy allows a non-invasive assessment of cell morphology, size variation of cell nuclei, blood vessel hyperplasia, or inflammatory reaction related aspects, which are important for assessing the state of a tissue(Benninger and Piston, 2013, Skala et al., 2005, Stanciu et al., 2014, Zipfel et al., 2003a). In SHG, two incident photons are combined into a single emitted photon with halved energy via a nonlinear process involving virtual states (Campagnola and Dong, 2011). One of the main applications of SHG for tissue characterization and diagnostics is imaging of collagen(Chen et al., 2012), which is the main structural protein in the extracellular matrix of animal tissues. Investigating collagen distribution with SHG enables a precise and non-invasive assessment of extracellular matrix modifications, which represent a hallmark of cancers(Bonnans et al., 2014, Lu et al., 2012), and of many other pathologies(Raines, 2000).

The usefulness of TPEF and SHG to characterize human skin has been demonstrated both ex-vivo (Paoli et al., 2009, Paoli et al., 2008) and in-vivo (Balu et al., 2015, Cicchi et al., 2014, Dimitrow et al., 2009, Koehler et al., 2011, Saager et al., 2015, Sun et al., 2017). Subsequently, MPM tomographs capable of providing virtual biopsies in-vivo have become available in many medical centers across the world(König, 2018). The utility of MPM techniques for characterizing skin, or other organs/parts of the human body, is manifold as they can (i) enhance our understanding of tissue anatomy and functionality, (ii) enable fast and accurate tissue characterization both ex-vivo and in-vivo. Because of these advantages, MPM is likely to soon become one of the central elements of in-vivo skin tissue characterization frameworks, while also representing a powerful tool to complement traditional diagnostics techniques such as immunohistochemistry or brightfield microscopy of H&E stained tissues. Recent advances in digital staining, where images taken by other modalities (including MPM) are transformed into virtual H&E images(Bocklitz et al., 2016, Borhani et al., 2019, Rivenson et al., 2019), facilitate the interpretation of MPM images by histopathologists, which we anticipate to massively boost the penetration of these techniques into the clinical practice.

In this work we perform MPM imaging of transversal tissue sections containing the dermoepidermal junction (DEJ)(Briggaman and Wheeler Jr, 1975), which separates the dermis and the epidermis, two morphologically distinct compartments that interact in several ways and at different levels to create, control, or restore tissue homeostasis. It is known that processes occurring near DEJ coordinate the growth and differentiation of the epidermis and are essential in demonstrating the complicated pathogenesis of epidermal tumors, irrespective of their benign or malignant nature. Our interest in characterizing this region is two-fold: (a) the DEJ lies at a depth accessible with MPM systems developed for clinical in-vivo applications(Breunig et al., 2018, Saager et al., 2015) (b) during epidermal carcinogenesis important changes take place in the DEJ, which can be linked to early hyperplastic and neoplastic phases. These changes can be classified by their appearance, their extent and their frequency, the most prominent ones being related to the destructive modifications occurring in the basal lamina when dealing with an invasive neoplastic proliferation (even in early stages). The first part of our experiment shows that many of these subtle changes, which are difficult to assess with conventional microscopy, are available with label-free MPM imaging of the DEJ.

The second part of our experiment finds motivation in the fact that manual evaluation approaches for histopathological image analysis and diagnostics are both time consuming and prone to errors (Brown, 2004, Chatterjee, 2014, Reid et al., 1988). To address this, we show that TPEF and SHG images collected on the DEJ can be analyzed by using Deep Learning (DL) (LeCun et al., 2015), to automatically and precisely distinguish between healthy and dysplastic skin tissues. Although our method is developed and tested on MPM datasets collected on fixed transversal tissue sections, it is on-the-fly translatable to in-vivo skin characterization assays for cancer screening/diagnostics based on clinically validated MPM tomographs capable to scan epithelial tissues in both horizontal and vertical directions (Breunig et al., 2018).

## RESULTS AND DISCUSSIONS

### MPM imaging of the DEJ for tissue state assessment

In Fig. 1 we present a set of MPM images (overlaid TPEF and SHG signals) of the DEJ, collected on normal and dysplastic epithelial tissues (transversal sections, see Methods). These are showed under a pseudo-coloring scheme: blue-color for collagen-rich tissues (providing contrast for SHG), red-color for autofluorescent tissue regions (probed by TPEF), and violet-color for co-localized SHG and TPEF signals. The displayed images demonstrate the utility of MPM signals collected on the DEJ in being helpful to a histopathologist in his task to assess the skin tissues state.

**Fig. 1.**
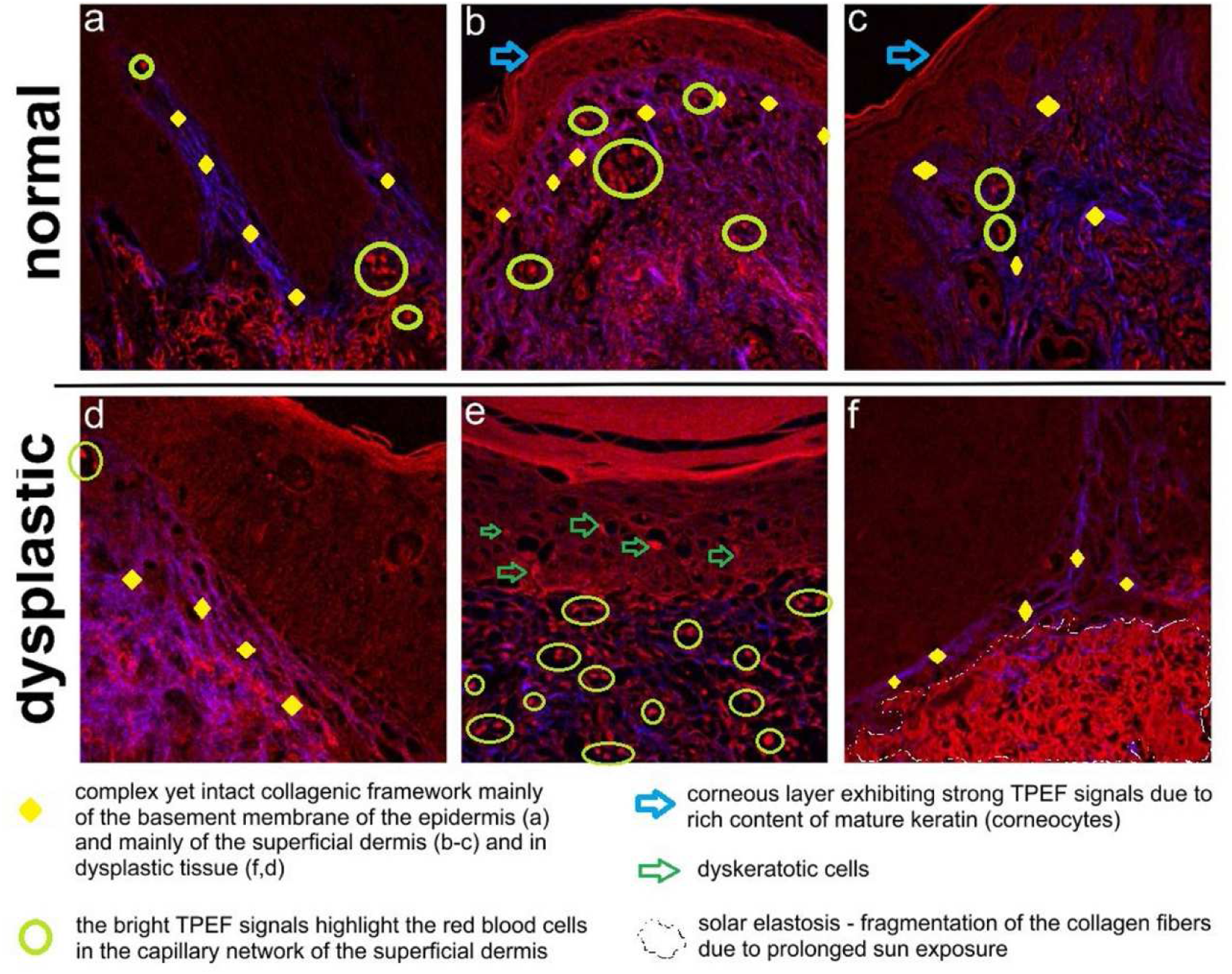
MPM images of the DEJ collected on normal and dysplastic epithelial tissues. Field of view: 250×250 µm^2^. (A version of these MPM images without marked elements is available as Supplementary Material, Fig. S1).

In Fig. 1a-c) we can observe a normal appearance of the DEJ, as the MPM images show a complex, yet intact, collagen framework mainly of the basement membrane of the epidermis (Fig. 1a) and of the superficial dermis (Fig. 1b,c), where the bright TPEF signals highlight the red blood cells in the capillary network of the superficial dermis. The delicate walls of the capillary are also highlighted by a continuous red line corresponding to TPEF signals. The epidermis exhibits homogeneous TPEF signals determined by cytokeratins inside the keratinocytes. We note that those signals have a monotonous cytoplasmic pattern in the squamous cells of the epidermis due to the keratin content. The keratinization is a dynamic process exhibiting a gradient of keratin content from the basal layer to the stratum corneum. Subsequently, increased TPEF signals in this layer correspond to a greater content of keratin. The corneous layer exhibits stronger TPEF emission having more mature keratin (corneocytes) – blue arrow in Fig. 1b,c); the epidermis has an overall honeycomb appearance. Fig. 1d-f) depict images of the DEJ in dysplastic tissues. In Fig. 1d,e) the strong TPEF signals in the papillary dermis originate from the hemoglobin in the red blood cells, which has been previously documented(Sun et al., 2015, Zheng et al., 2011). The presence of cells with abnormal individual keratinization (dyskeratotic cells) is shown by TPEF signals similar of those of the red blood cells which are marked with green arrows in Fig. 1e). TPEF signals also indirectly outline the nuclear contour of the squamous cells, which is important for assessing their state (the irregular nuclear contour is an important feature of neoplastic lesions, in situ or malignant). The increased nuclear ratio, dyskeratotic cells and parakeratosis are common features of actinic keratosis. In the case of the basement membrane the collagen framework is still visible and suggests an in-situ lesion, but the rest of the collagen framework of the papillary dermis has a more fragmented pattern suggesting a degenerative process (prolonged solar exposure). Fig. 1f) is representative for the MPM images collected on tissues affected by actinic keratosis, showing the usefulness of this technique to highlight the degeneration and fragmentation of the collagen fibers due to solar elastosis (area marked by dotted line).

A landmark study (Barsky et al., 1983) showed that the basement membrane, one of the main components allowing the identification of the DEJ in the case of healthy and dysplastic tissues, is lost in the case of invasive tumors of the skin. The MPM images collected on malignant tissues, Fig. 2, containing relevant borders of such tumors, are in accordance with this previous study, the DEJ no longer being visible. These images contain nonetheless features that allow a histopathologist to assess the tissue state, and tumor invasion patterns. For example, In Fig. 2a) we can observe a region corresponding to normal epidermis (left-area marked by dotted line) demonstrated by monotonous cytoplasmic TPEF signals, and to the basement membrane nearby, easily observable based on strong SHG signals. On the right, we can observe similar cytoplasmic signals in a moderate differentiated squamous cell carcinoma (SCC); few keratin pearls (green circles in Fig 2a-c) or dyskeratotic cells are seen (blue arrows in Fig. 2a-c). We observe here that the large irregular nuclei are indirectly highlighted by TPEF (yellow dotted line in Fig. 2a-c). This front of invasion shows a pushing borders invasion pattern on the remaining structures. The collagen in the remaining dermis is fragmented (shown by SHG dot-like blue signals and marked by orange diamonds in Fig. 2a-c). Interestingly, solar elastosis has a strong, almost homogenous red signal in TPEF suggesting that this process promotes the presence of endogenous chromophores in regions affected by this condition. Fig. 2a-c) are also representative for SCC. In these three images we can easily observe the large dyskeratotic cells (blue arrows) with strong TPEF signals. Large, individual or grouped cells with squamous differentiation are found in the papillary dermis, admixed with red blood cells (with very strong TPEF signals – marked by red circles) and lymphocytes (rounded small cells with a dimmer appearance – marked by blue circles); Irregular, parakeratotic pearls can also be seen (green circles). The collagen framework, highlighted by blue signals from SHG, is partially destroyed, suggesting an invasive lesion.

**Fig. 2.**
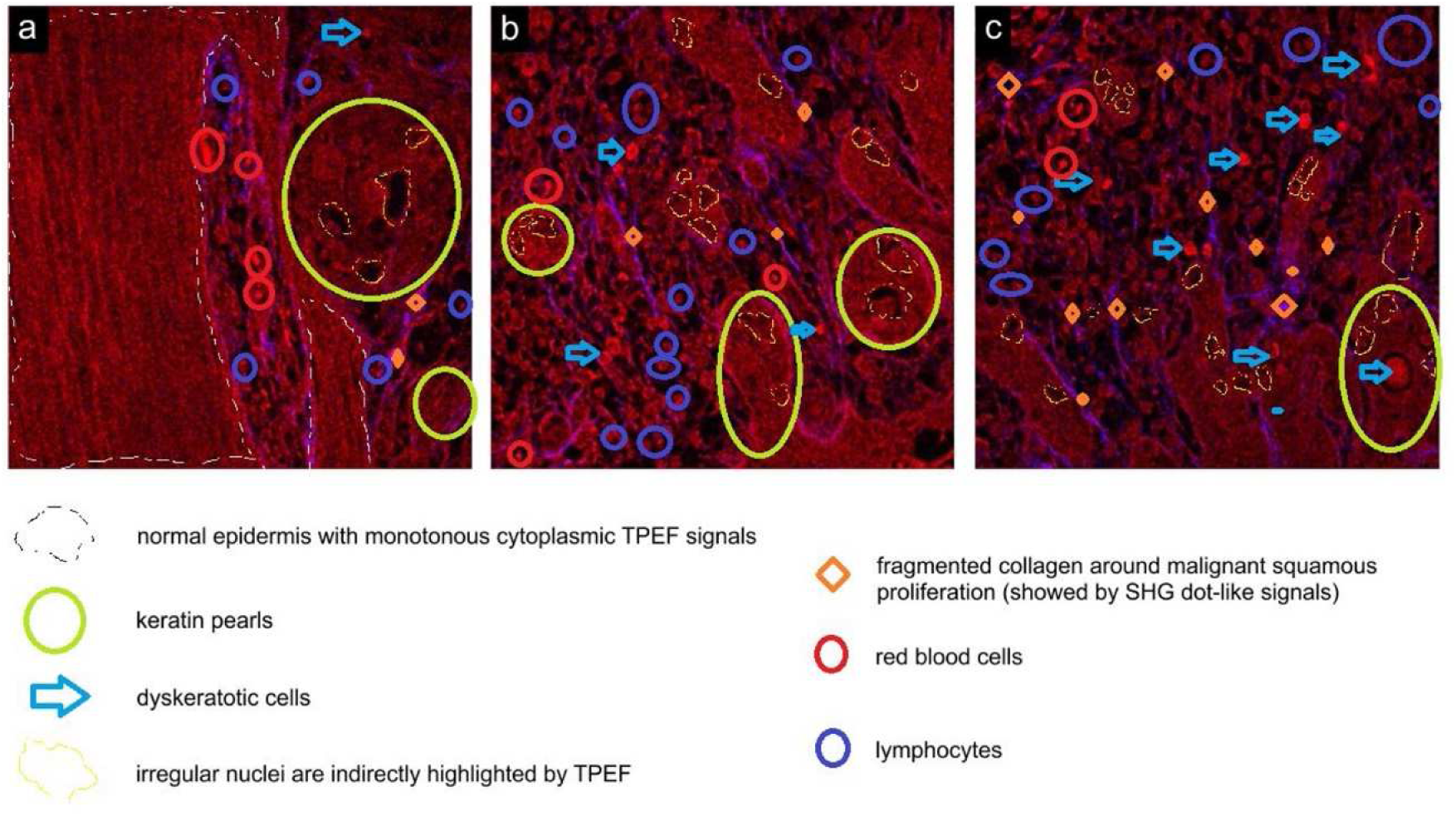
MPM images collected on malignant epithelial tissues on DEJ related regions. Field of view: 250×250 µm^2^. (A version of these MPM images without marked elements is available as Supplementary Material, Fig. S1).

Following our analysis of the MPM images collected on epithelial tissues, it can be observed that the DEJ region can be easily identified in the case of healthy and dysplastic tissues.

In the case of malignant tissues, components of the DEJ are lost, this region being compromised and no longer identifiable. MPM imaging of transversal tissue sections containing the DEJ holds thus potential for screening/diagnostics, in fixed, fresh and in-vivo samples, allowing a histopathologist to differentiate healthy from dysplastic (unlabeled) tissues. Such utility is of great importance especially with respect to in-vivo assays, since identifying dysplastic lesions is often difficult with non-invasive methods, and patients are reluctant to allow the physician to resort to excisional biopsy when they are not convinced of the risk. Furthermore, such non-invasive in-vivo screening/diagnostic methods based on transversal (xz scanning direction) MPM imaging of the DEJ would represent a key tool for patients who cannot be subjected to excisional biopsy without the risk for complications(Abhishek and Khunger, 2015), e.g. patients suffering of hemophilia(Chapin et al., 2017), or in patients where cutaneous excision may result in a defect that is difficult to correct by plastic surgery(Bayat et al., 2003).

### Automated identification of healthy and dysplastic tissues with MPM and Deep Learning

The first part of our experiment showed that the DEJ is easily identified in MPM images collected on transversal sections of healthy and dysplastic tissues and provides important cues for a histopathologist to assess the tissue state. In malignant tissues the DEJ is compromised and hence not identifiable. Considering these, we hypothesized that an important utility of DEJ investigation with MPM systems dedicated to clinical imaging, e.g. (Balu et al., 2015, Koehler et al., 2011, König, 2008, Weinigel et al., 2014), would refer to potential assays that aim to screen patients with dysplastic modifications of the skin, which are difficult to implement with traditional non-invasive modalities. To further explore this utility, in the second part of our experiment we implemented and evaluated a DL method that augments the potential of MPM imaging of the DEJ, aiming to achieve automated and precise classification of tissues either as healthy or dysplastic.

The employed DL image classification method was inspired from a recent work(Huttunen et al., 2018), and dealt with a total of 358 MPM images from healthy (*n* = 14) and dysplastic (*n* = 14) unstained tissue sections, collected to contain the DEJ (see Methods). Images were randomly divided into validation (70%) and training sets (30%), the latter being augmented with the two strategies: (i) by reflecting the original images horizontally and vertically, and (ii) by repeatably blurring these horizontal and vertical reflections by using a five-layer Gaussian image pyramid, (see Methods). The first augmentation strategy resulted in 1000 training images, while application of both yielded a set consisting of 5000 images. The test set always consisted of 108 images. This DL classification experiment was repeated 25 times, each time using a random selection of the validation/training images.

Previous work showed that better classification accuracy of MPM images of (ovarian) tissues is achieved by exploiting images with merged (summed) TPEF and SHG signals, compared to addressing solely TPEF or SHG images (Huttunen et al., 2018). Here, we have extended the number of MPM signals for automated tissue classification, by additionally including in our evaluation framework the metabolic redox ratio (REDOX), which is calculated based on the TPEF emission originating from FAD and NADH molecules in the tissue(Georgakoudi and Quinn, 2012, Skala et al., 2007). We have thus experimented DL classification of MPM derived images representing the following summed-up signals: a) SHG+TPEF, b) SHG+TPEF+REDOX, c) TPEF+REDOX, d) SHG+REDOX. The resulting mean sensitivities, specificities and accuracies along with their standard deviations are shown in Fig. 3.

**Fig. 3.**
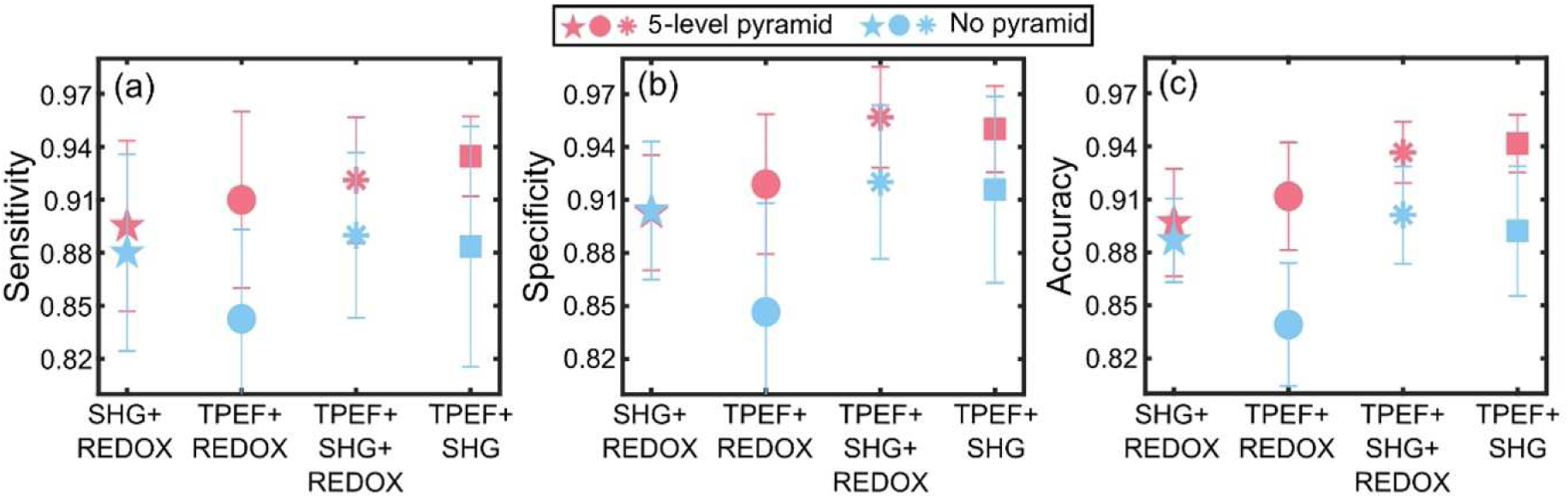
Calculated classification sensitivity (a), specificity (b) and accuracy (c) with the error bars corresponding to the respective standard deviations. (a)-(c) Classification performance is on average improved ∼4 % when the data augmentation includes also a 5-level Gaussian blur pyramid (red markers), compared to data augmentation not utilizing the pyramid (blue markers). (a)–(c) Highest classification sensitivity (93.5 ± 2.3%), specificity (95.0 ± 2.4%) and accuracy (94.2 ± 1.6%) are achieved by using combined TPEF+SHG images for training.

The classification performance of the trained network (GoogLeNet) for all four different MPM signal permutations was in general outstanding (∼90% or better). The best classification sensitivity (93.5 ± 2.3%, see Fig. 3a), specificity (95.0 ± 2.4%, see Fig. 3b) and accuracy (94.2 ± 1.6%, see Fig. 3c) were achieved using combined TPEF+SHG images. Use of “TPEF+SHG+REDOX” images resulted in best classification specificity (95.7 ± 2.8%, see Fig. 3b) alongside with excellent sensitivity (92.1 ± 3.6%) and accuracy (93.7 ± 1.7%). The classification performance was on average improved ∼4 % when the image pyramid scheme was employed, compared to the case when data augmentation was done only with horizontal and vertical reflections of the original images. While the number of training images (5000 vs 1000) could also explain this improvement, we believe that the main underlying reason refers to the fact that different layers of a Gaussian image pyramid approximate images of the original objects collected at different scales(Florack et al., 1992, Sporring et al., 2013) (the reason why we adopted this data augmentation strategy). Therefore, by including layers of the image pyramid in the training process it is as if we expose the network to images collected at different magnifications, conferring thus scale invariance to the classification framework. The advantage of scale invariance for the performance of DL classification of MPM images collected on epithelial tissues derives from the fact that the morphology of skin varies with age(Branchet et al., 1991), anatomical site(Gambichler et al., 2006, Huzaira et al., 2001) and other factors. Convolving laser-scanning microscopy images (in our case the MPM images used for training) with a Gaussian filter is also known to suppress noise(Van Kempen et al., 1997), which might as well have contributed to the observed classification performance increase.

### Conclusions

In this work we have focused our attention on MPM imaging of transversal tissue sections containing the DEJ, a region of the skin which is known to harbor important processes and modifications that are relevant with respect to the pathogenesis of epidermal tumors. We showed that MPM images contain features that are easy to interpret, which allow assessing the integrity of the DEJ, and differentiating healthy from dysplastic tissues. We regard this as being of great medical interest because compromised DEJ structures are a major hallmark of cancer progression and invasiveness and identifying these at a very early stage of the disease in a non-invasive manner compatible with in-vivo deployment has great importance for timely implementing the appropriate therapeutic strategies. Secondly, we have shown that MPM images of the DEJ, besides being easily interpreted by a trained expert, can also be automatically classified with DL either as healthy or dysplastic. To this end, we have demonstrated a DL approach based on the GoogLeNet network, which provides real-time image classification, with sensitivity, specificity and accuracy all exceeding 90%.

The demonstrated methodology for automated classification of MPM data sets collected on the DEJ can be on-the-fly transferred to *in-vivo* screening assays based on clinically validated multiphoton tomographs, which are already available in many institutions worldwide. In such assays, a target region of the patient’s skin should be scanned with an MPM tomograph in xz direction (transversally), and once the DEJ is visible, an MPM image containing it could be instantly classified as healthy or dysplastic with the demonstrated DL strategy (or adapted variants). Overall, our results show that MPM and DL are likely to play a huge role in the forthcoming years in terms of speeding up and improving the current methodologies used in skin cancer screening and diagnosis.

## METHODS

### Image acquisition and processing

Combined SHG and TPEF imaging (to which we refer throughout the paper as MPM imaging) was performed using an upright Leica TCS-SP confocal laser-scanning microscope modified for nonlinear optical imaging. We used for excitation a Ti:Sapphire laser (Chameleon Vision II, Coherent) tuned at 860 nm, with ∼140 fs pulses and a repetition rate of 80 MHz. A 40X magnification and 0.75 numerical aperture objective was used for focusing the laser beam on the sample and for collecting the backward-generated MPM signals (Fig. 4, representation adapted from (Stanciu et al., 2017)). The average power reaching the sample plane was kept below 15 mW. Images were acquired with a linear laser beam polarization obtained by using a polarization stage generator (PSG) comprised of an achromatic quarter-wave plate (AQWP05M-980, Thorlabs) and an achromatic half-wave plate (AHWP05M-980, Thorlabs), mounted in motorized rotation stages (PRM1/MZ8, Thorlabs) and placed in the laser beam path before the microscope. Three different input polarizations at 0°, 60° and 120° were used. The spectral detection available with the Leica TCS SP microscope allowed us to collect three channels simultaneously: the SHG channel (425 – 435 nm) and two TPEF channels – 440 – 490 nm corresponding to intrinsic fluorescence of reduced pyridine nucleotides (NAD(P)H) and 510 – 600 nm detecting the flavin adenine dinucleotide (FAD) fluorescence. The SHG/TPEF images were the Kalman average of four consecutive frames. The final SHG image was obtained by averaging the three SHG images acquired at different laser beam polarizations, resulting in a polarization independent SHG image(Gao et al., 2006). Image processing was performed using FIJI(Schindelin et al., 2012), and consisted of applying a 0.5 radius mean filter to reduce noise. In addition, contrast was automatically enhanced, and a 0.7 gamma correction was applied in order to enhance the visibility of low intensity collagen fibers in the SHG images. The TPEF images corresponding to the NAD(P)H and FAD configurations were processed similarly, and the final TPEF image was formed by averaging the two separate TPEF NAD(P)H and FAD images. A composite RGB image was obtained by inserting the processed SHG and TPEF images into the blue and red channels, respectively.

Brightfield microscopy (BM) images of H&E stained samples were collected using a Leica DM 3000 LED brightfield microscope, equipped with an MC 190 HD camera. A HC PL Fluotar 5x/0.15 ∞/-/OFN25/C objective was used to record overlapping image tiles that were stitched together to form large mosaics representing the entire sample slides. The sample regions that were imaged with MPM were also imaged with BM at high magnification (50X).

**Fig. 4.**
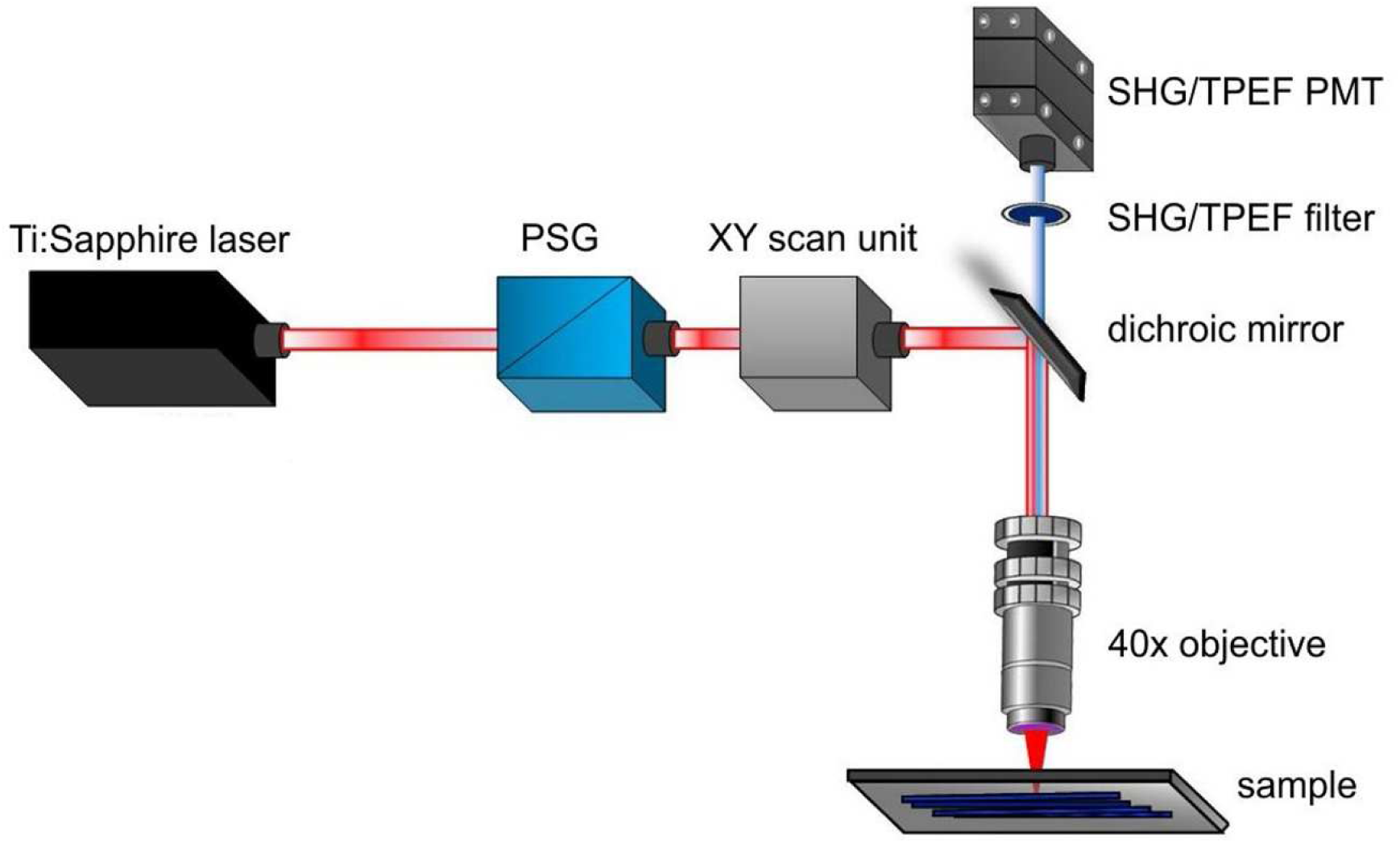
Configuration of the MPM imaging setup.

### Sample preparation and imaging

The skin tissue samples used in this experiment consist in (i) lesions typical to SCC, which has been regarded as an ideal prototype for lesions with a malignant/invasive character, (ii) lesions that are considered premalignant/non-invasive/“in situ”, e.g. from patients diagnosed with actinic keratosis or Bowen disease. Normal skin-tissue fragments have also been included in the study and were obtained from healthy regions close to the resection margins of the considered malignant/premalignant lesions. For each of the three investigated tissue categories we have imaged 14 pairs of histological slides, each corresponding to a distinct case. To obtain a pair of histological slides, from a formaldehyde-fixed paraffin-embedded histological block, two skin tissue sections were consecutively cut; one was left unstained for MPM imaging, while the other was stained with H&E for conventional histopathology. Using BM mosaics collected on this latter, trained histopathologists marked the positions of the DEJ, and MPM images were collected at random sites across the marked DEJ on the unstained sample pair. The regions imaged with MPM were also imaged with BM at high magnification (50X) for ground-truth. The imaging framework is presented in Fig. 5.

**Fig. 5.**
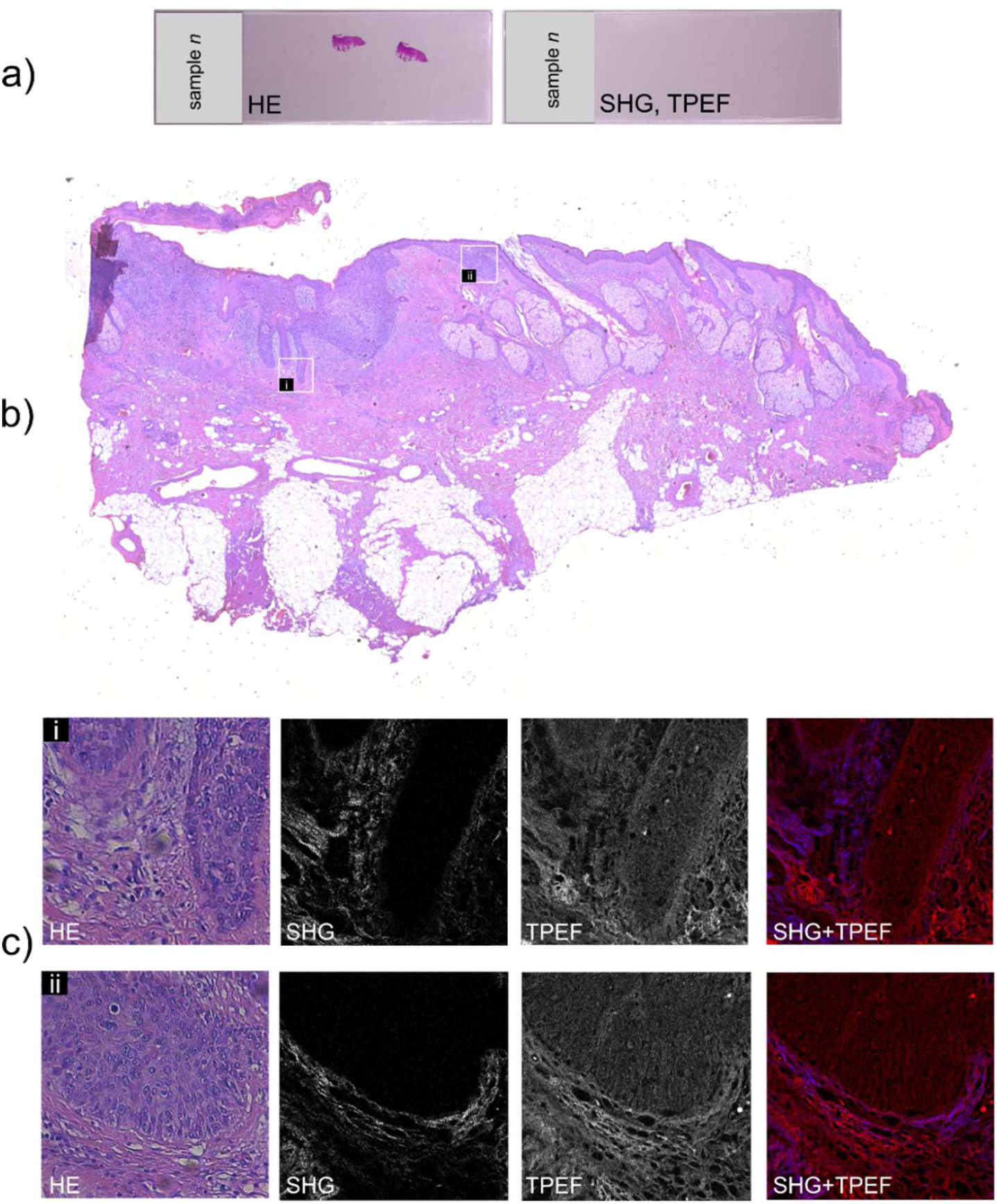
Schematic demonstration of the employed imaging protocol. A) Representative photograph of two consecutively cut tissue fragments, the first stained with H&E, and the second left unstained. B) Large mosaic depicting the entire histology slide assembled by stitching overlapping BM images (5X obj.) C) Example of MPM images (40X obj.) collected on the unstained samples at random positions across the DEJ. All acquired MPM images were registered to high magnification BM images (50X obj.) collected on the corresponding regions of the H&E stained sample, in order to re-confirm that they indeed depict the DEJ, which is captured transversally.

### MPM image classification with Deep Learning

For classifying the MPM images we employed a pre-trained convolutional neural network (GoogLeNet), originally trained to perform 1000-fold multi-class classification of images by using a database consisting of ∼1.2 million annotated images(Deng et al., 2009, Szegedy et al., 2015). Typically, very large data sets are needed to train networks from scratch and to overcome problems related to overfitting(Krizhevsky et al., 2012). To address this, we used a pre-trained network(Deng et al., 2009, Szegedy et al., 2015) which alleviates the cumbersome problem of generating large scale MPM training datasets(Huttunen et al., 2018). This way, we were able to use a relatively small data set of MPM images of the DEJ to fine-train the network to perform binary classification of target MPM images. As annotated training data was available, a supervised learning scheme was employed(Erhan et al., 2010). Prior to the fine-training, we replaced the final classification layers of the original network (layers after ‘*pool5*’) with new fully connected (FC), softmax and classification output layers in order to perform binary classification of MPM images. The data workflow of the approach is illustrated in Fig. 6.

**Fig. 6.**
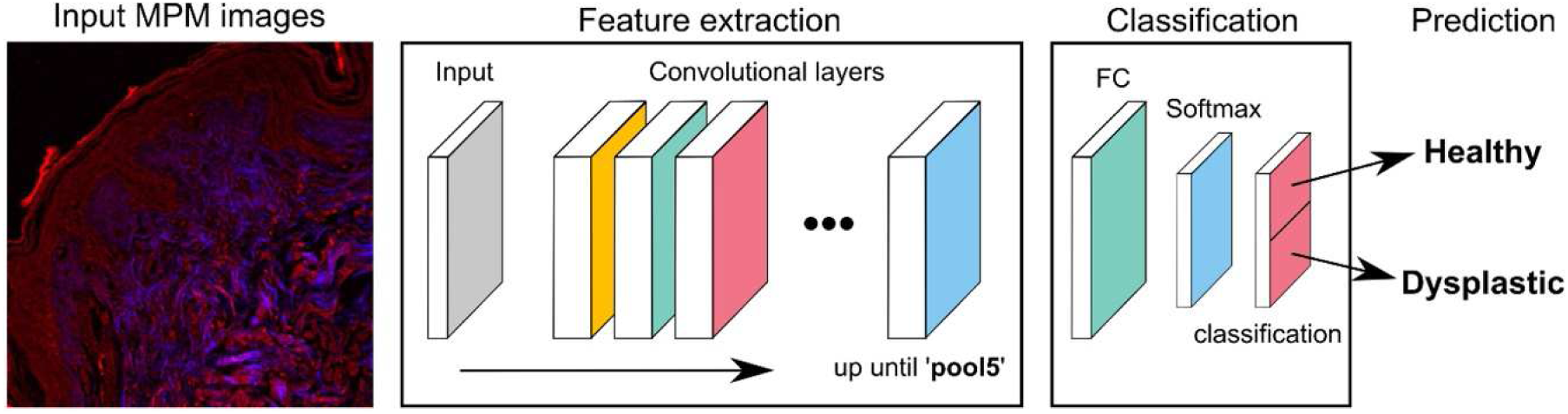
Convolutional Neural Network based workflow for binary classification of MPM images. The input images are fed to the pre-trained network (GoogLeNet) that first performs feature extraction effectively transforming the data into a more optimal representation. Subsequent image classification is performed by the FC, Softmax and classification layers that, contrary to the convolutional layers, are trained from scratch during the fine-training process.

To further address overfitting problems raised by the relatively small training data set, (Krizhevsky et al., 2012), we employed data augmentation, which is known to help in this matter(Wang and Perez, 2017). We augmented MPM training data in two ways: (i) by casting horizontal and vertical reflections (a commonly met strategy), and (ii) with a novel strategy that combines horizontal/vertical reflections with the image pyramid in the Gaussian-Scale Space(Adelson et al., 1984). Here, each layer of the image pyramid (5 in total: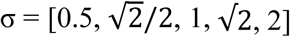) was horizontally and vertically reflected. The effect of the Gaussian blur pyramid scheme on an MPM image of the DEJ is illustrated in Supplementary Material, Fig. S2.

## Acknowledgments

Part of the work of A. Dumitru, M. Costache, R. Hristu and S.G. Stanciu was financially supported by the Romanian Executive Agency for Higher Education, Research, Development and Innovation Funding (UEFISCDI), under Grant PN-III-P2-2.1-PED-2016-1252 MICAND. S.G. Stanciu ad R. Hristu were also supported by the UEFISCDI grant PN-III-P1-1.1-TE-2016-2147 (CORIMAG) and by the ATTRACT project funded by the EC under Grant Agreement 777222, (via grant HARMOPLUS). S.G. Stanciu also acknowledges the support of the COST Action CA15124 NEUBIAS which facilitated interactions relevant for this work, and the support of the NVIDIA Corporation, through their academic hardware grant awarded to support his research on Deep Learning. Work with the Chameleon Vision II (Coherent) fs laser, was possible due to European Regional Development Fund through Competitiveness Operational Program 2014-2020, Priority axis 1, Project No. P_36_611, MySMIS code 107066, Innovative Technologies for Materials Quality Assurance in Health, Energy and Environmental - Center for Innovative Manufacturing Solutions of Smart Biomaterials and Biomedical Surfaces – INOVABIOMED. M. J. Huttunen acknowledges the support from the Academy of Finland (308596) and the Flagship of Photonics Research and Innovation (PREIN) funded by the Academy of Finland (320165). The authors thank Mr. Tiberiu Totu (UPB) and Ms. Roxana Buga (UPB) for their help with acquisition of BM mosaics.

## Competing interests

The authors declare no competing interests.

## Authors contributions

SGS, RH, AD and MC designed the experiments dealing with MPM imaging of the DEJ. RH collected the MPM data sets and part of the BM datasets and further on dealt with their digital processing for improved visualization. AD and MC selected relevant cases to be included in the performed studies and prepared the samples. AD annotated the BM images for consistent acquisition of MPM datasets. SGS, AD and MC analyzed the MPM data sets collected on the DEJ. MJH and SGS designed the experiments dealing with automated classification of MPM datasets via Deep Learning. MJH implemented the work on Deep Leaning, which was done at Tampere University. MJH, SGS and MC analyzed the results achieved on automated MPM data classification with Deep Learning. All authors wrote and reviewed the manuscript.

