## Supplemental Figures for "Investigating and Assessing the Dermoepidermal Junction with Multiphoton Microscopy and Deep Learning"

### SUPPLEMENTARY FIGURES

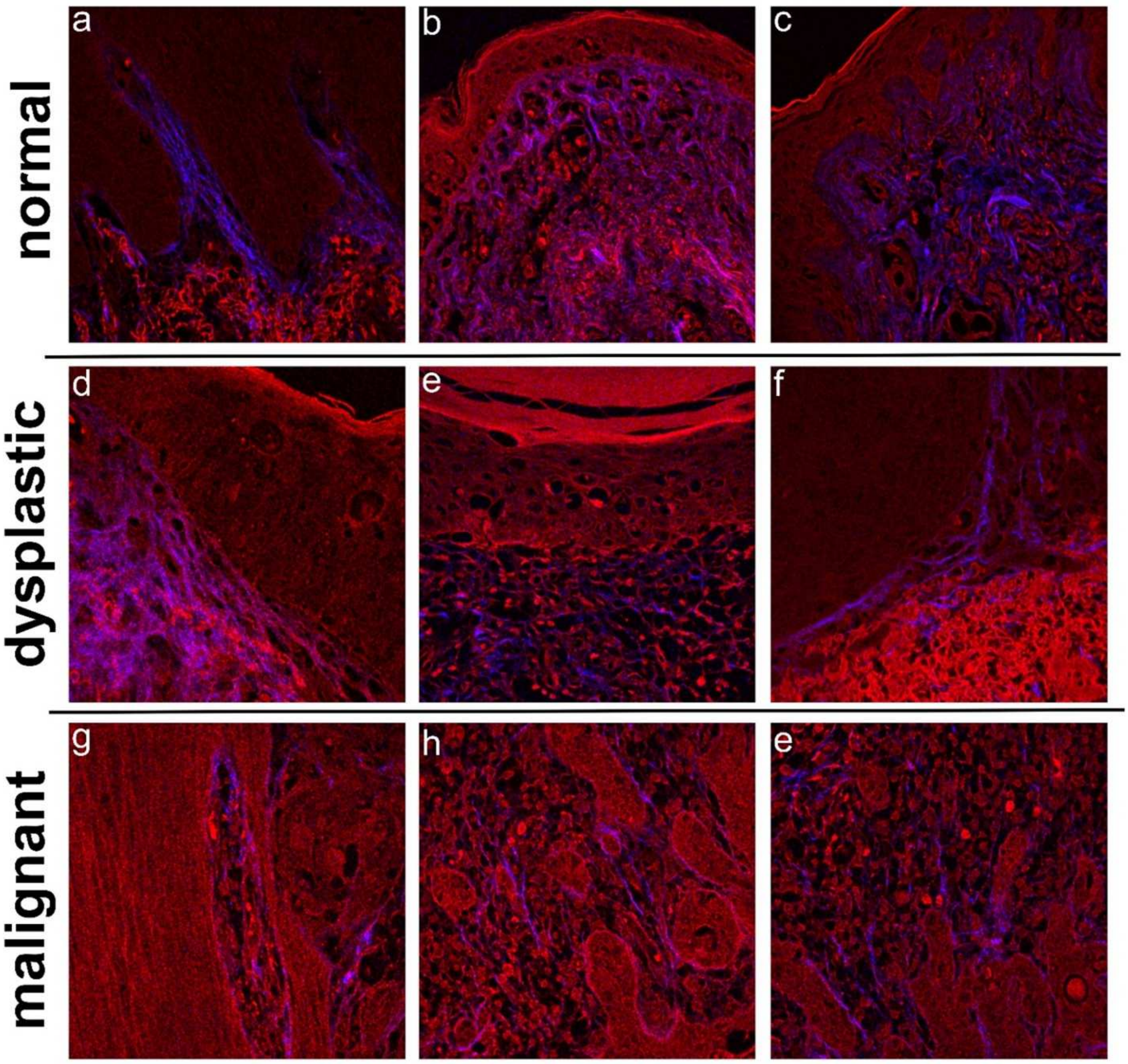


**Fig. S1**. MPM images (TPEF in red color, SHG in blue color) collected on normal, dysplastic and malignant epithelial tissues. The MPM images of healthy and dysplastic tissues were acquired to contain the dermo-epidermal junction. In malignant tissues the dermo-epidermal junction is compromised, and hence not observable in the displayed MPM images. Field of view: 250x250 $\mu m^2$.

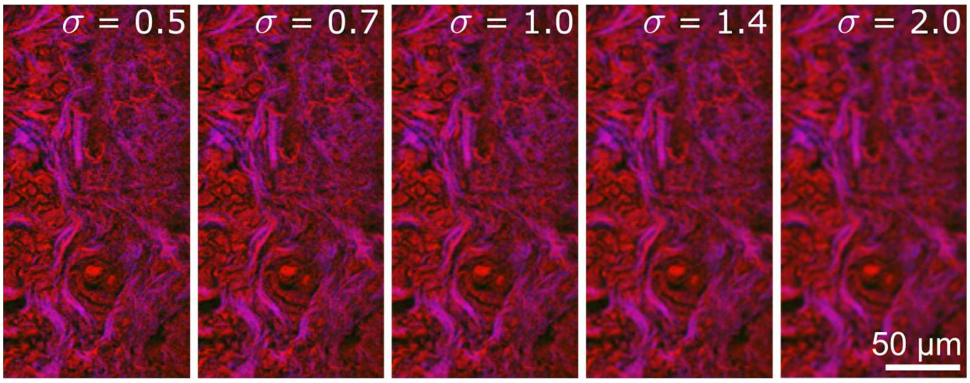


**Fig. S2.** Representative augmented MPM training images by performing 5-level Gaussian blur using standard deviations σ = 0.5, $\sqrt{2}/2$, 1, $\sqrt{2}$ and 2.
